# A bacterial effector blocks SUMOylation by steric occlusion of UBC9 via arginine-GlcNAcylation

**DOI:** 10.64898/2026.03.06.710069

**Authors:** Zhongrui Ma, Yaoyao Zhu, Yanheng Zhao, Xiangyu Wang, Huajie Zhang, Bingqing Li

## Abstract

The post-translational modifier SUMO is crucial for host antimicrobial defense. Here, we discover that *Salmonella* employs a unique strategy to dismantle the host SUMOylation machinery. *Salmonella* infection robustly inhibits global SUMOylation, particularly SUMO2/3 conjugation, without altering the expression of SUMO-cycle enzymes. Genetic screening identified the T3SS-2 effector SseK1 as both necessary and sufficient for this suppression. SseK1, an arginine-GlcNAcyltransferase, directly binds and specifically modifies the SUMO E2-conjugating enzyme UBC9 at arginine 17 (R17). Structural and biophysical analyses revealed that GlcNAcylation at R17 sterically hinders UBC9’s interaction with SUMO, thereby inactivating the entire SUMOylation cascade. A unique C-terminal lid domain (ARHVQ motif) in SseK1 confers substrate specificity for UBC9, representing an evolutionary innovation within *Salmonella* genus. Quantitative SUMOylome profiling demonstrated that SseK1-mediated UBC9 inactivation reprograms the host SUMOylation landscape, impairing the modification of key immune regulators such as MyD88, Hspa8 and destabilizing proteins like PDCD4. Consequently, this mechanism is critical for *Salmonella* intracellular survival and systemic virulence in mice. Our study unveils a unique bacterial strategy to dismantle the host SUMOylation machinery via precise enzymatic inactivation of its central hub.

## Introduction

Post-translational modifications (PTMs) are fundamental orchestrators of the innate immune response^1^. Among these, SUMOylation—the covalent attachment of small ubiquitin-like modifiers (SUMOs) to lysine residues—regulates the activity, stability, and subcellular localization of the target proteins^2, 3, 4^. This modification is particularly critical in the modulation of Toll-like receptor (TLR) pathways^5^, inflammasome activation^6^, DNA-damage responses^7^, and transcriptional control^8^. By dynamically altering SUMOylation landscape or SUMOylome, the hosts enable a rapid defense response against diverse bacterial and viral challenges^9, 10^.

To counteract these antimicrobial defenses, numerous bacterial pathogens have evolved sophisticated strategies to subvert host SUMOylation machinery^11, 12, 13^.The most prevalent mechanism involves the targeted degradation of SUMO-cycle enzymes. For instance, *Listeria monocytogenes* secretes listeriolysin O (LLO) to trigger the degradation of the SUMO E2-conjugating enzyme UBC9^14^; *Staphylococcus warneri* utilizes the toxin Warnericin RK to deplete both UBC9 and the SUMO E1-activating enzyme SAE1/2^15^; and *Shigella* exploits calcium/calpain-mediated cleavage of SAE2^16^. Beyond degradation, other strategies have emerged: *Rhizobia* effector NopD exhibits intrinsic deSUMOylation activity^17^, while *Klebsiella pneumoniae* promotes the cytosolic accumulation of the deSUMOylase SENP2 by inhibiting its ubiquitination and subsequent degradation^18^.

*Salmonella enterica*, a major intracellular pathogen causing both localized gastroenteritis and systemic disease, has also been implicated in suppressing host SUMOylation. Previous studies suggested that *Salmonella* achieves this via microRNA-mediated silencing of *Ube2i* (encoding UBC9) and *Pias1* (a SUMO E3 ligase)^19^, or through c-Fos-mediated transcriptional repression of their promoters^20^. However, transcriptional silencing alone is insufficient to explain the rapid and profound loss of SUMOylation observed as early as 2-4 hours post-infection^19^. Given that UBC9 and PIAS1 possess predicted half-lives of ∼30 hours (analyzed by ProtParam), even a complete arrest of transcription would not cause a collapse of the existing protein pool within such a short timeframe. Meanwhile, paradoxically, our transcriptomic and proteomic analyses demonstrate that SUMO-cycle enzymes remain stable during both the early and late stages of *Salmonella* infection in cell culture and mouse models. These observations suggest that *Salmonella* employs a previously unrecognized, non-degradative strategy to rapidly subvert host SUMOylation.

In this study, we demonstrate that the *Salmonella* Type III Secretion System 2 (T3SS-2) is indispensable for the inhibition of host SUMOylation. Through a systematic screen, we identified the T3SS-2 effector SseK1 as the factor responsible for this blockade. SseK1 is an arginine-GlcNAcyltransferase previously known to modify death domain-containing proteins^21, 22, 23^; however, its role in hijacking the SUMOylation machinery has not been reported.

Using yeast two-hybrid screening, we identified UBC9 as the primary target of SseK1 in macrophages. Distinct from pathogens that deplete UBC9, *Salmonella* utilizes SseK1 to directly catalyze the GlcNAcylation of UBC9 at residue R17. This modification sterically disrupts the interaction between UBC9 and its SUMO substrates, effectively paralyzing host SUMOylation. We further show that SseK1 recruits UBC9 via a unique C-terminal lid domain, which is absent in all other known arginine-GlcNAcyltransferases, highlighting a highly specialized evolutionary adaptation of *Salmonella*.

Finally, the global impact of *Salmonella* infection on the host SUMOylome remains largely uncharacterized. By leveraging quantitative SUMO-proteomics, we identified 303 SUMOylated proteins during *Salmonella* infection. Herein, SseK1 selectively suppresses the SUMOylation of several critical immune regulators, including MyD88, Hspa8, and PDCD4. Cell-based and mouse infection models further confirm that this SseK1-mediated SUMOylome reprogramming is essential for *Salmonella* pathogenesis. Collectively, our findings uncover a novel enzymatic strategy of “modification-mediated inactivation” used by *Salmonella* to subvert host immunity and define the SseK1-UBC9 axis as a battleground in host-pathogen interactions.

## Results

### *Salmonella* subverts host SUMOylation without altering SUMO-cycle enzyme expression

To investigate the impact of *Salmonella* infection on host SUMOylation, we first infected RAW264.7 macrophages with *Salmonella enterica* serovar Typhimurium (STM) strain ATCC 14028S. Immunofluorescence microscopy at 4 hrs post-infection (hpi) revealed a profound reduction in global SUMOylation levels, which were mainly attributable to decreased SUMO2/3 conjugation, compared with uninfected cells (Fig. 1a).

**Fig. 1.**
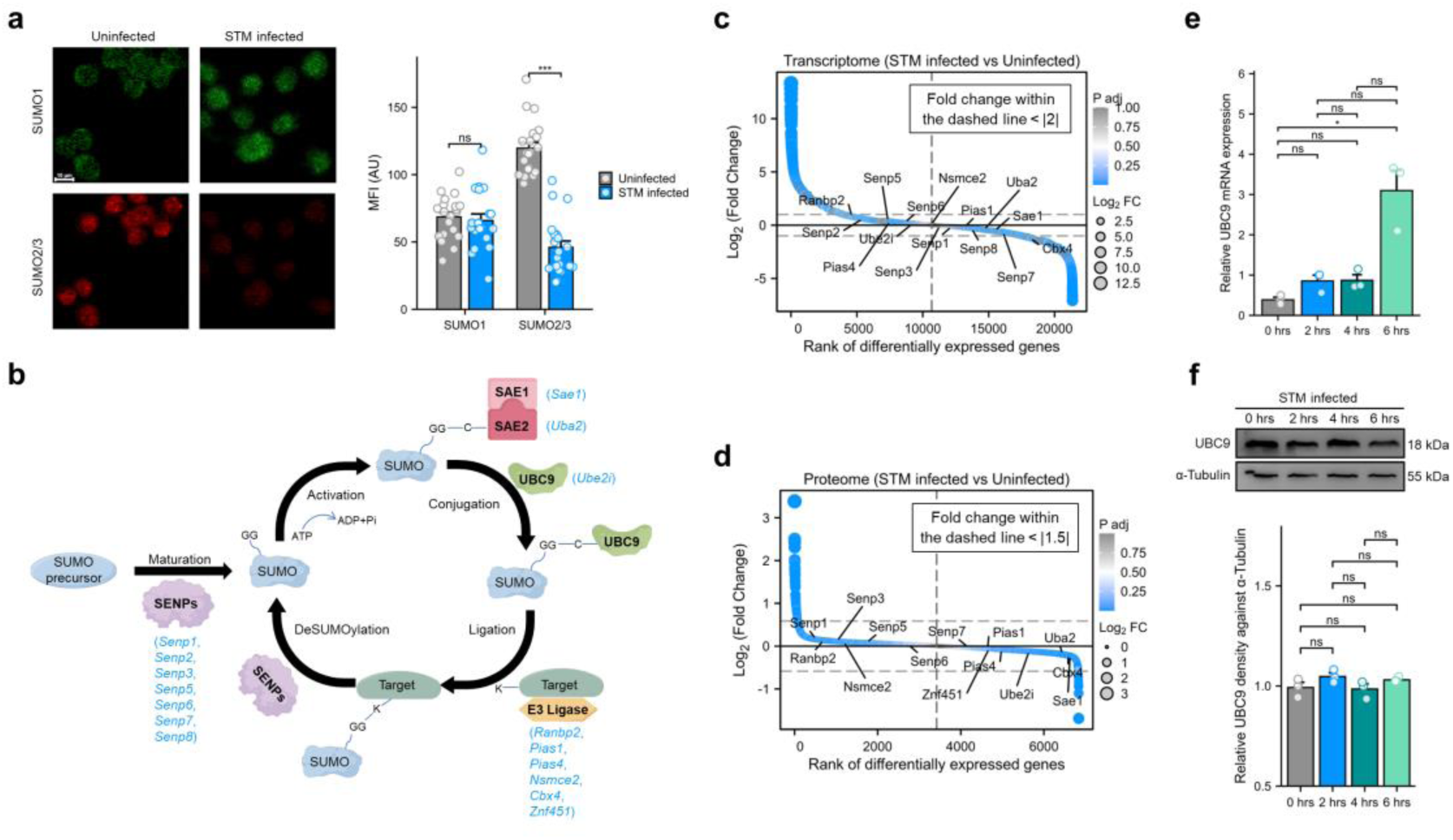
*Salmonella* subverts host SUMOylation without altering SUMO-cycle enzyme expression. **a** Immunofluorescence images and quantification (n = 20) of SUMO1 (green) and SUMO2/3 (red) conjugation in RAW264.7 cells with or without STM 14028S infection (4 hpi). Scale bar, 10 µm. **b** Core SUMO cycle enzymes and their corresponding primary genes (blue). **c, d** Transcriptomic and proteomic analyses of SUMO cycle enzyme expression in RAW264.7 cells infected with STM 14028S vs uninfected controls (4 hpi; n = 3). **e** UBC9 mRNA expression in RAW264.7 cells during STM 14028S infection (0-6 hpi; n = 3). **f** Immunoblot analysis and quantification of UBC9 protein levels in STM 14028S-infected RAW264.7 cells (0-6 hpi; n = 3). *P < 0.05; ***P < 0.001; ns, not significant.

Intriguingly, despite this robust phenotypic loss of SUMOylation, the mRNA abundance of core SUMO-cycle enzymes—including SAE1/2, UBC9, ligases, and deSUMOylases (Fig. 1b)—remained largely unchanged (fold change < 2) following STM infection (Fig. 1c). Protein levels of these enzymes were likewise stable (fold change < 1.5) (Fig. 1d). To ensure that this stability was not an artifact of our specific cell model or time point, we cross-referenced our results with multiple independent, publicly available datasets. Analysis of transcriptomic profiles from STM-infected Caco-2 cells (24 hpi)^24^ or mouse intestinal tissues (4 dpi)^25^, and *Salmonella* Enteritidis-infected rat colonic mucosa (3 dpi) or Caco-2 cells (2 hpi)^26^ consistently showed that SUMO-cycle gene expression remains unperturbed across various stages of infection and host species (Supplementary Fig. 1). Given its indispensable role in the SUMOylation cascade, we further scrutinized UBC9 expression via qPCR and immunoblotting (Fig. 1e, f). Both UBC9 mRNA and protein levels remained constant during the first 6 hrs of infection. Notably, a modest increase in UBC9 mRNA was detected at 6 hpi (Fig. 1e), which we hypothesize may represent a compensatory host feedback response to the impaired SUMOylation machinery. Collectively, these data demonstrate that *Salmonella* infection robustly subverts host SUMOylation, particularly SUMO2/3 conjugation, through a mechanism that is independent of SUMO-cycle enzyme degradation or transcriptional silencing.

### T3SS-2 effector SseK1 is essential for *Salmonella*-mediated subversion of host SUMOylation

To elucidate the mechanism by which *Salmonella* subverts host SUMOylation, we interrogated the temporal dynamics of host SUMOylation during STM infection. In RAW264.7 cells, SUMO1 conjugation remained largely unaffected within the first 6 hrs, whereas SUMO2/3 conjugation began to decline precipitously starting at 2 hpi (Fig. 2a). The kinetics of this decline coincided with the established activation profile of the *Salmonella* T3SS-2, which typically begins to translocate effectors into the host cytoplasm around 2 hpi. This synchronicity led us to hypothesize that T3SS-2 serves as the primary apparatus for subverting host SUMOylation. Supporting this, infection with a T3SS-2-deficient mutant (Δ*ssaV*) failed to induce the depletion of SUMO2/3 conjugates at 4 hpi, with SUMOylation levels remaining indistinguishable from those of uninfected controls (Fig. 2b).

**Fig. 2.**
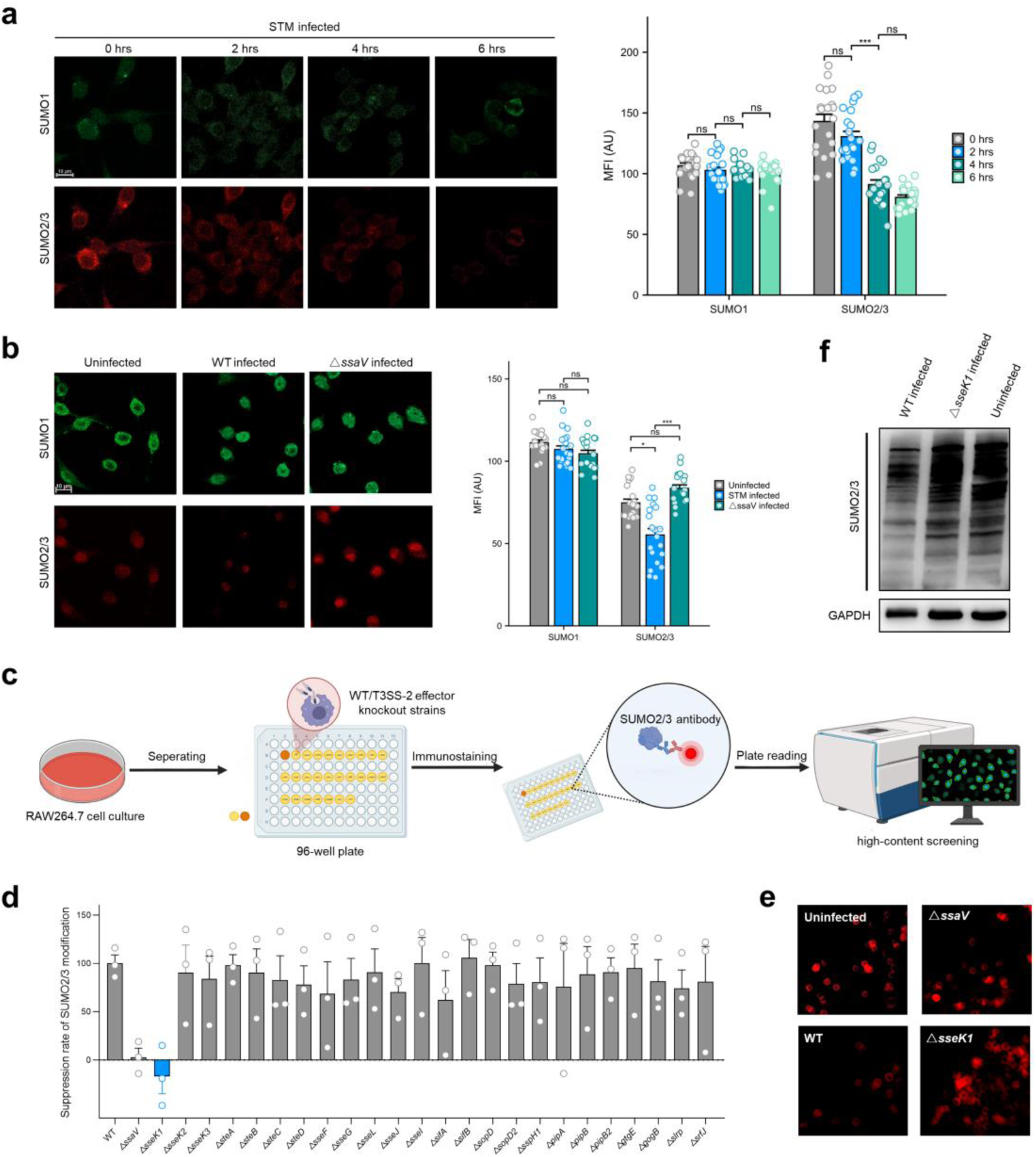
*Salmonella* subverts host SUMOylation through T3SS-2 effector SseK1. **a** Immunofluorescence images and quantification (n = 20) of SUMO1 (green) and SUMO2/3 (red) conjugation in RAW264.7 cells infected with STM 14028S over a 0-6 hrs time course. Scale bar, 10 µm. **b** Immunofluorescence images and quantification (n = 20) of SUMO1 (green) and SUMO2/3 (red) conjugation in RAW264.7 cells infected with STM WT or Δ*ssaV* or left uninfected (4 hpi). Scale bar, 10 µm. **c** Schematic of the high-content screening (HCS) workflow used to identify T3SS-2 effector(s) required for subversion of host SUMOylation. **d** SUMO2/3 suppression rates in RAW264.7 cells infected with STM WT or T3SS-2 effector knockout strains (4 hpi). For each sample, fluorescence intensity (FI) was measured across three fields (20 cells per field) to calculate the mean FI (MFI). Suppression rate = (MFI_uninfected - FI_test) / (MFI_uninfected - MFI_WT). **e** Representative immunofluorescence images of SUMO2/3 conjugation in RAW264.7 cells infected with STM WT, Δ*ssaV*, Δ*sseK1*, or left uninfected (4 hpi). **f** Immunoblot analysis of SUMO2/3 conjugates in RAW264.7 cells infected with STM WT or Δ*sseK1* or left uninfected (4 hpi). *P < 0.05; ***P < 0.001; ns, not significant.

Given that T3SS-2 delivers a diverse repertoire of effector proteins to manipulate host physiology, we sought to identify the specific effector(s) responsible for this SUMOylation blockade. We utilized a systematic library of 24 individual T3SS-2 effector knockout mutants and performed a high-content screening (HCS) based on SUMO2/3 fluorescence intensity (Fig. 2c). Among the candidates tested, only the Δ*sseK1* strain exhibited a significant loss of SUMO2/3-suppressive activity, phenocopying the Δ*ssaV* mutant (Fig. 2d, e). Immunoblot analysis further confirmed that SseK1 is strictly required for the reduction of SUMO2/3 conjugates during infection (Fig. 2f). Taken together, these data pinpoint SseK1 as the definitive T3SS-2 effector employed by *Salmonella* to subvert the host SUMOylation landscape.

### SUMO E2-conjugating enzyme UBC9 is identified as principal host target of SseK1

To identify the host substrates targeted by SseK1, we performed an unbiased yeast two-hybrid (Y2H) screen using SseK1 as bait against a RAW264.7 macrophage cDNA library (Fig. 3a). This screen yielded 18 positive clones (Supplementary Table 1), among which four independent hits were mapped to *Ube2i*, the gene encoding the SUMO E2-conjugating enzyme UBC9 (Fig. 3b, c). Given that UBC9 is the sole and indispensable E2 conjugase in the SUMOylation cascade, we prioritized it as the primary candidate for further investigation.

**Fig. 3.**
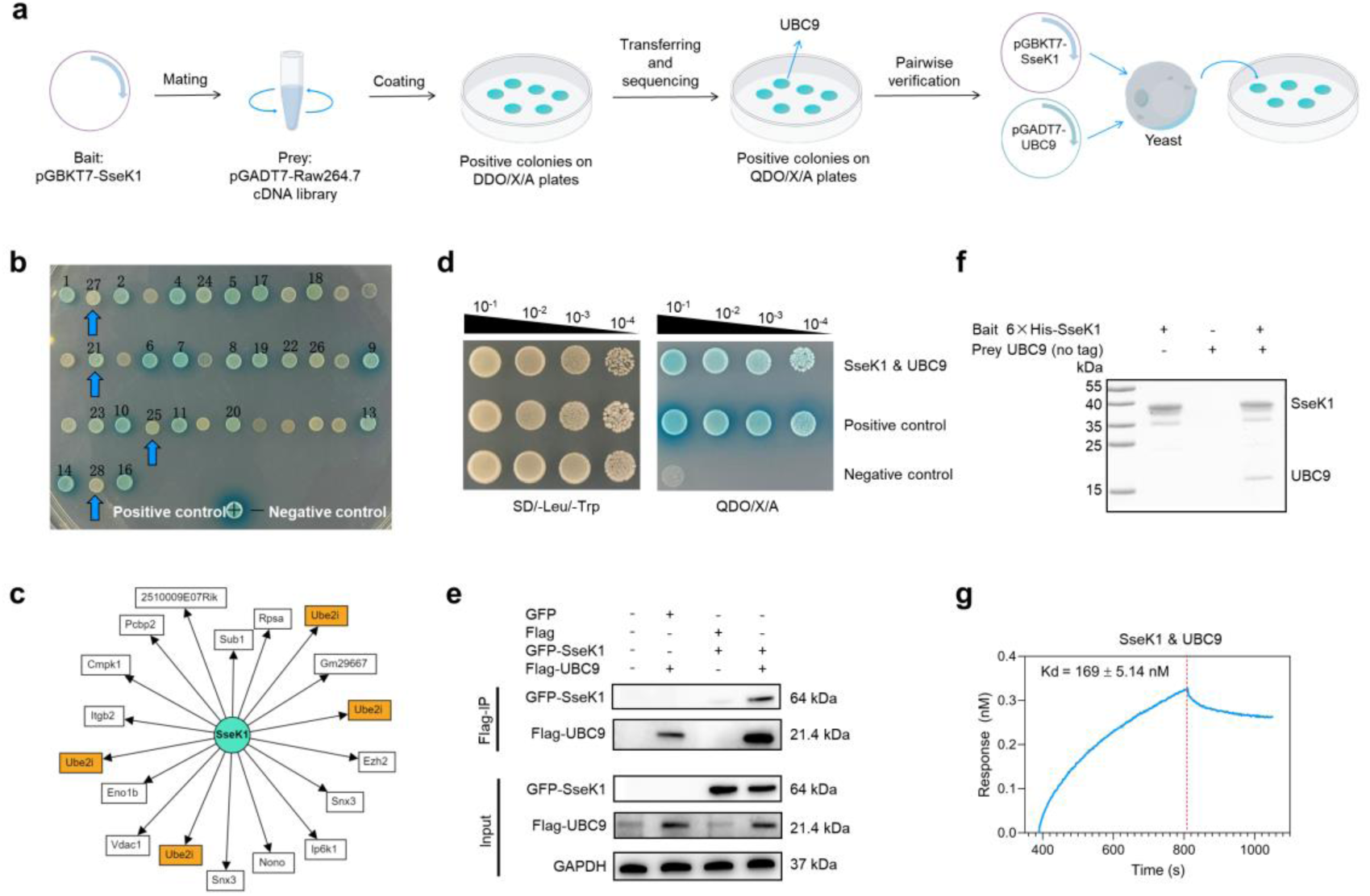
SseK1 interacts with UBC9. **a** Schematic of the Y2H screening workflow and pairwise interaction validation strategy. **b** Representative positive colonies on selective plates; four colonies (blue arrows) correspond to *Ube2i* (encoding UBC9). **c** Sequencing analysis of positive clones showing that four independent clones encode *Ube2i* (orange). **d** Pairwise interaction assay validating the interaction between SseK1 and UBC9. **e** Co-IP in HEK293T cells confirming the interaction between SseK1 and UBC9. **f** *In vitro* pull-down assay demonstrating direct binding between purified SseK1 and UBC9. **g** BLI analysis showing the binding kinetics between SseK1 and UBC9 *in vitro*.

Pairwise Y2H assays confirmed a direct interaction between SseK1 and UBC9 in yeast (Fig. 3d). We next validated this interaction in mammalian cells. Co-immunoprecipitation (Co-IP) in HEK293T cells demonstrated that SseK1 robustly associates with UBC9 in a cellular context (Fig. 3e). To determine whether this interaction is direct and independent of other host factors, we purified recombinant SseK1 and UBC9 proteins (Supplementary Fig. 2) for *in vitro* biochemical analysis. Pull-down assays confirmed the formation of a stable SseK1-UBC9 complex (Fig. 3f). Furthermore, we quantified the binding kinetics using bio-layer interferometry (BLI), which revealed a high binding affinity with a dissociation constant kd of 169 ± 5.14 nM (Fig. 3g). Collectively, these findings establish that SseK1 directly and high-affinity binds to UBC9, identifying it as a principal host target during *Salmonella* infection.

### SseK1 catalyzes arginine-GlcNAcylation of UBC9 at R17 during *Salmonella* infection

SseK1 is a specialized glycosyltransferase that catalyzes the transfer of N-acetylglucosamine (GlcNAc) onto the arginine (R) residues of target substrates. Given our observation that SseK1 physically associates with UBC9, we investigated whether UBC9 serves as a direct enzymatic substrate. We first monitored the modification status of Flag-tagged UBC9 in HEK293T cells during *Salmonella* infection. While infection with STM WT triggered robust arginine-GlcNAcylation of UBC9, this modification was entirely abolished in cells infected with the Δ*sseK1* strain (Fig. 4a), demonstrating that SseK1 is required for UBC9 modification *in vivo*. To determine whether SseK1 directly catalyzes this reaction, we performed *in vitro* enzymatic assays using purified recombinant proteins. SseK1 efficiently GlcNAcylated both bacterially expressed 6×His-UBC9 (Fig. 4b) and Flag-UBC9 isolated from mammalian cells (Supplementary Fig. 3), indicating that the modification is directly mediated by SseK1 and independent of additional host factors.

**Fig. 4.**
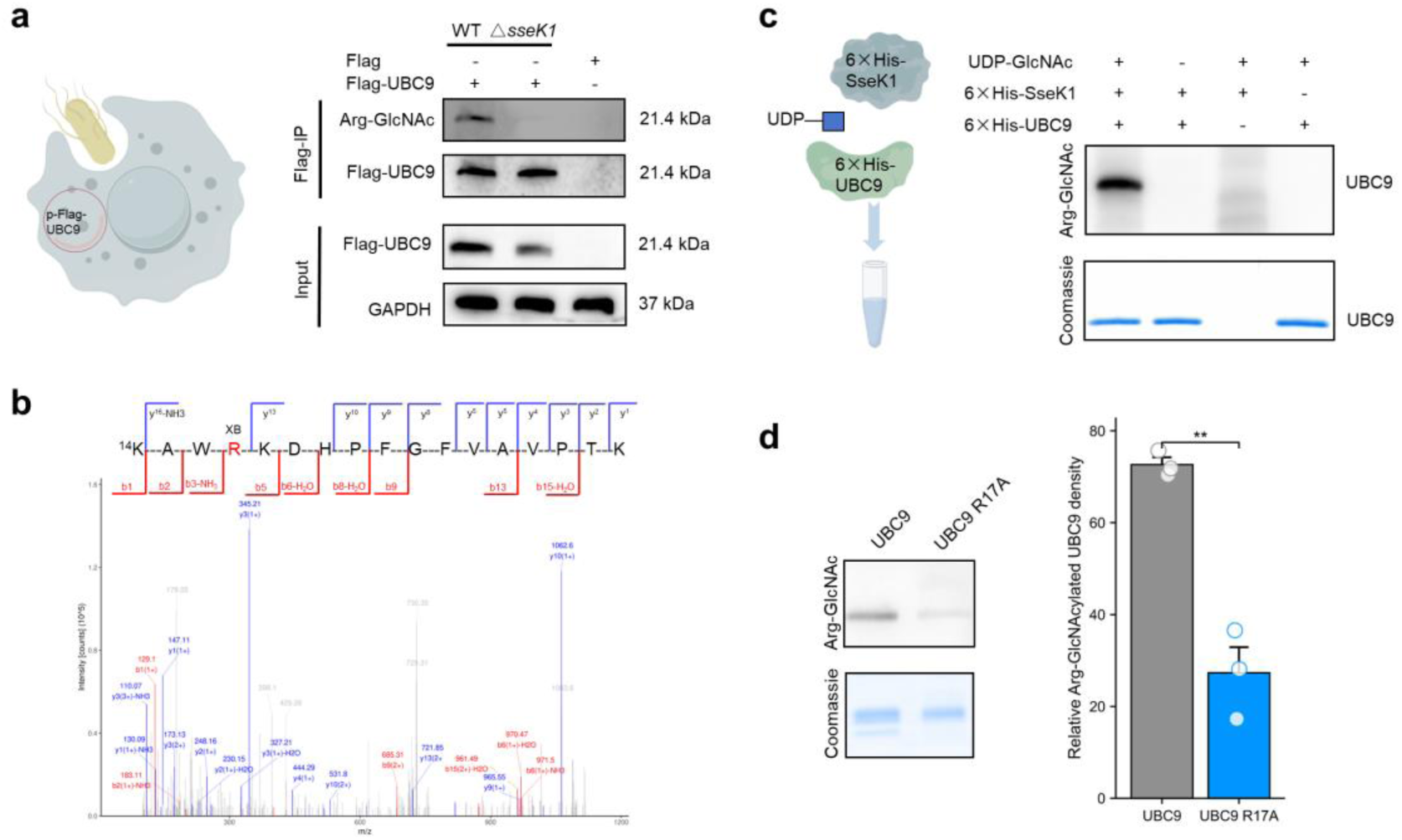
SseK1 catalyzes arginine-GlcNAcylation of UBC9. **a** HEK293T cells expressing Flag-UBC9 were infected with STM WT or Δ*sseK1*, followed by immunoprecipitation and immunoblot analysis of UBC9 Arg-GlcNAcylation. **b** *In vitro* enzymatic assays showing Arg-GlcNAcylation of bacterially expressed 6×His-UBC9, detected by immunoblotting. **c** MS/MS spectrum of Arg-GlcNAcylated Flag-UBC9 peptides. Corresponding b and y ions are indicated along the peptide sequence above the spectrum. **d** Immunoblot analysis and quantification (normalized to Coomassie-stained protein, n = 3) of Arg-GlcNAcylation of UBC9 and UBC9 R17A. **P < 0.01.

To pinpoint the modified residue(s), we subjected arginine-GlcNAcylated Flag-UBC9, which was purified from STM WT-infected cells, to mass spectrometry analysis. The mass spectra identified R17 of UBC9 as the specific GlcNAcylation site (Fig. 4c). To validate this finding, we generated a UBC9 mutant with an R17-to-alanine (R17A) substitution. In contrast to native UBC9, the R17A mutant exhibited a profound reduction in arginine-GlcNAcylation in *in vitro* enzymatic assays (Fig. 4d). Collectively, these data establish that SseK1 functions as an arginine-GlcNAcyltransferase that specifically targets the R17 residue of UBC9 during *Salmonella* infection.

### UBC9 R17 GlcNAcylation sterically hinders SUMO binding

UBC9 serves as the central E2-conjugating enzyme in the SUMOylation cascade, and its non-covalent recruitment of SUMO is a prerequisite for subsequent thioester formation and substrate transfer. Previous studies have implicated the R17 residue of UBC9 as a critical determinant of this SUMO-binding interface^27^. To explore the structural implications of R17 modification, we analyzed the crystal structure of the UBC9–SUMO1 complex (PDB: 2UYZ) and a predicted model of the UBC9–SUMO2 interaction. Our analysis positioned R17 squarely within the binding interface, where it facilitates essential hydrogen bonding with the SUMO moiety (Fig. 5a, b). We therefore hypothesized that the addition of a bulky GlcNAc moiety at this site would sterically or electrostatically disrupt these intermolecular contacts, thereby attenuating UBC9 function. To test this hypothesis, we utilized a bacterial co-expression system to produce homogeneously Arg-GlcNAcylated UBC9 (Fig. 5c), confirming the modification site at R17 via mass spectrometry (Supplementary Fig. 4). In parallel, SUMO1 and SUMO2 proteins were individually purified from bacterial expression systems (Fig. 5d). We then quantified the impact of this modification on binding kinetics using MicroScale Thermophoresis (MST). Compared to unmodified UBC9, the R17-GlcNAcylated form completely abolished the binding affinity for SUMO2 (showing no detectable binding signal), whereas its interaction with SUMO1 was attenuated but still retained a measurable response (Fig. 5e, f). Notably, the more profound inhibitory effect on SUMO2 binding provides a mechanistic basis for our observation that SseK1 preferentially suppresses the SUMO2/3-conjugated pool during *Salmonella* infection. Collectively, these biophysical data demonstrate that SseK1-mediated GlcNAcylation effectively decouples UBC9 from its SUMO partners, leading to a systemic collapse of the host SUMOylation machinery.

**Fig. 5.**
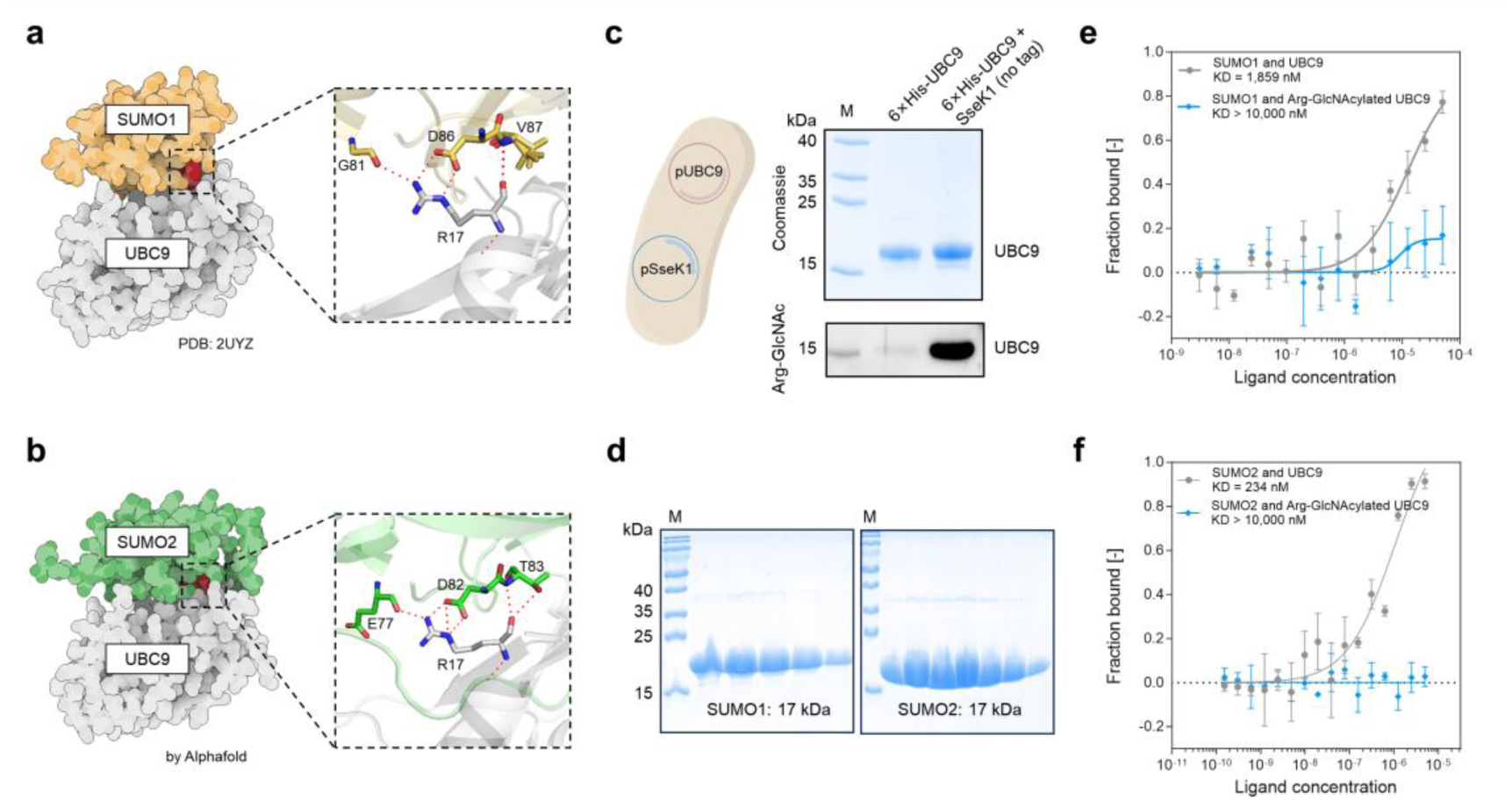
UBC9 R17 GlcNAcylation attenuates UBC9-SUMO binding affinity. **a** Crystal structure of the UBC9-SUMO1 complex (PDB: 2UYZ). **b** Predicted structure of the UBC9-SUMO2 complex generated using AlphaFold3. Enlarged views of the UBC9-SUMO1 and UBC9-SUMO2 interfaces are highlighted in black boxes. Residues forming hydrogen bonds with UBC9 R17 are shown in stick representation. **c** Purification of 6×His-UBC9 from *E. coli* BL21(DE3) co-expressing pSseK1 and pUBC9, followed by immunoblot analysis of UBC9 Arg-GlcNAcylation. **d** Purification of SUMO1 and SUMO2 proteins from *E. coli* BL21(DE3) harboring pSUMO1 or pSUMO2, respectively. M, molecular weight marker. **e, f** MST analysis of the binding affinities between SUMO1 or SUMO2 and unmodified UBC9 or Arg-GlcNAcylated UBC9.

### A unique C-terminal lid domain dictates SseK1 substrate specificity for UBC9 recruitment

Despite the high degree of homology between the *Salmonella* glycosyltransferases SseK1, SseK2, and SseK3, we observed a striking functional divergence: only SseK1 could suppress host SUMOylation (Fig. 2d and Supplementary Fig. 5). This specificity was recapitulated in cell-free enzymatic assays, where SseK2 and SseK3 failed to catalyze the arginine-GlcNAcylation of UBC9 (Fig. 6a). To identify the structural basis for this selectivity, we performed in silico modeling using AlphaFold3. While the three paralogs share a conserved global architecture, SseK1 possesses a unique C-terminal lid domain (Fig. 6b). Structural docking of the SseK1-UBC9 complex predicted that residues H334 and V335 within this lid domain form critical hydrogen bonds with T35 and N31 of UBC9, respectively, effectively acting as a molecular clamp to stabilize the enzyme-substrate interface (Fig. 6c).

**Fig. 6.**
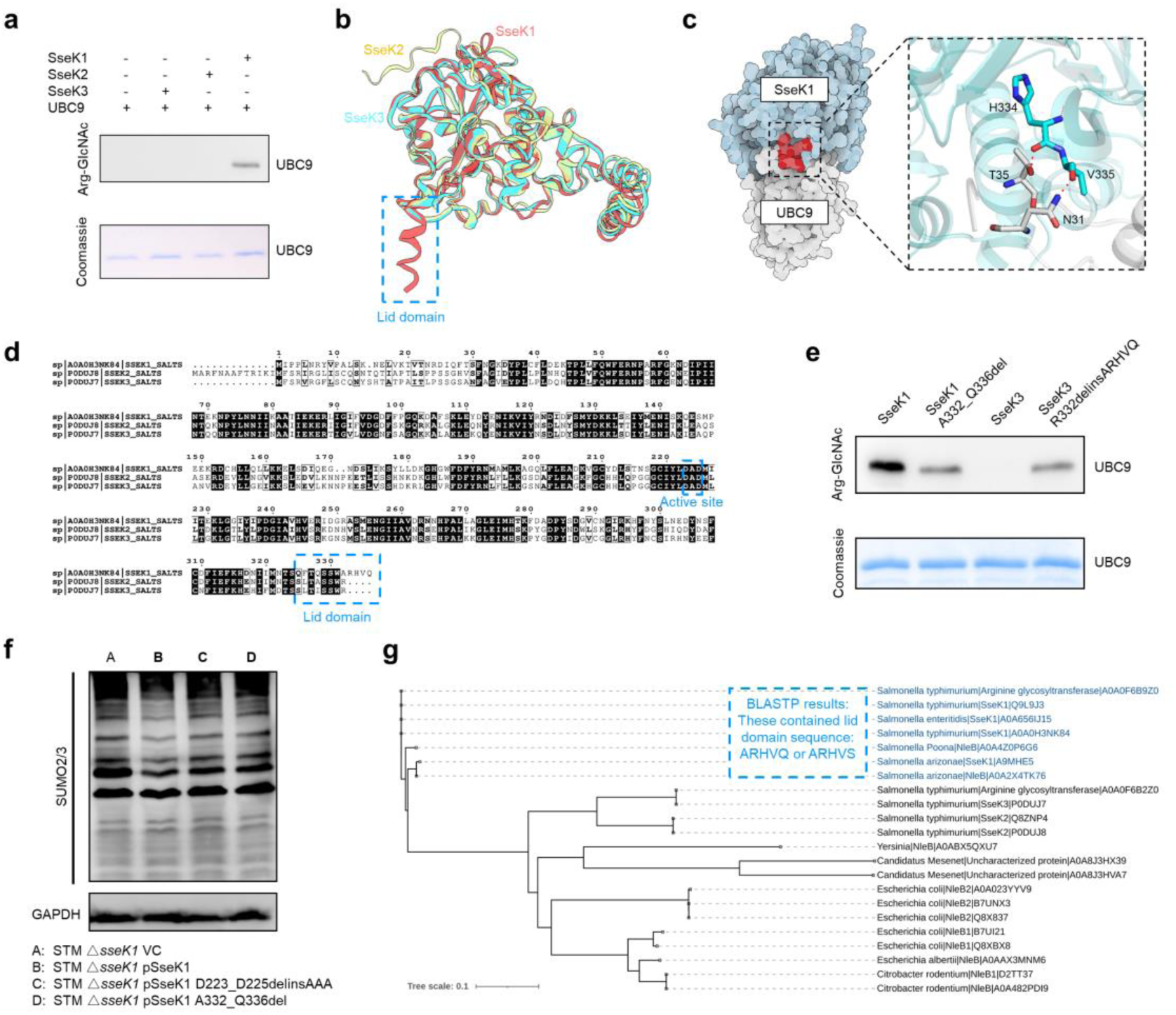
SseK1 recognizes UBC9 through a unique lid domain. **a** Immunoblot analysis of UBC9 Arg-GlcNAcylation mediated by SseK1, SseK2, or SseK3. **b** AlphaFold-predicted structures of SseK1 (aa 29-336, red), SseK2 (aa 29-348, yellow), and SseK3 (aa 29-335, green), shown with structural alignment using PyMOL. **c** AlphaFold-modeled structure of the SseK1-UBC9 complex. The enlarged view of the lid-domain region is boxed in black. Residues in the SseK1 lid domain forming hydrogen bonds with UBC9 are shown in stick representation. **d** Sequence alignment of SseK1, SseK2, and SseK3 generated using ESPript 3.0. **e** Immunoblot analysis of UBC9 Arg-GlcNAcylation mediated by SseK1, SseK1 A332_Q336del, SseK3, and SseK3 R332delinsARHVQ. **f** Immunoblot analysis of SUMO2/3 conjugation in RAW264.7 cells infected with STM Δ*sseK1*, STM Δ*sseK1* complemented with SseK1, or STM Δ*sseK1* complemented with either SseK1 D223_D225delinsAAA or SseK1 A332_Q336del (4 hpi). **g** Phylogenetic analysis of *Salmonella* Typhimurium SseK1 homologs. Protein sequences homologous to SseK1 (UniProt accession: A0A0H3NK84) were identified using BLASTP analysis against the UniProtKB reference proteomes and Swiss-Prot databases. Sequences with an E-value < 0.05 were selected for phylogenetic analysis. The phylogenetic tree was constructed and visualized using the Interactive Tree of Life (iTOL) online tool.

Sequence alignment identified a characteristic ARHVQ motif within this lid domain as a unique feature of SseK1 (Fig. 6d). To test the functional significance of this element, we performed domain-swapping and deletion experiments. Targeted deletion of the lid domain (SseK1 A332_Q336del) abolished SseK1-mediated UBC9 arginine-GlcNAcylation (Fig. 6e). Remarkably, grafting the ARHVQ motif onto SseK3 (SseK3 R332delinsARHVQ) was sufficient to confer partial GlcNAcylation activity toward UBC9, demonstrating that the lid domain is a primary determinant of substrate recruitment (Fig. 6e). Consistent with these findings, wild-type SseK1 restored SUMOylation blockade in Δ*sseK1*-infected RAW264.7 cells, whereas lid-deficient (A332_Q336del) and catalytically inactive (D223_D225delinsAAA) mutants did not (Fig. 6f).

To explore the evolutionary context of this mechanism, we performed phylogenetic analysis across diverse bacterial arginine-GlcNAcyltransferases. We found that SseK1 and a subset of its homologs form a distinct lineage defined by the presence of the lid-domain motif (ARHVQ or the variant ARHVS) (Fig. 6g). Intriguingly, this sequence appears to be an evolutionary innovation exclusive to the *Salmonella* genus, identified across multiple serovars including Typhimurium, Enteritidis, Poona, and Arizonae. In contrast, SseK1-like proteins from other enteric pathogens, such as *Escherichia coli*, *Citrobacter rodentium*, and *Yersinia spp.*, lacked this specific motif and did not group within this lineage. Collectively, these results reveal that *Salmonella* has evolved a unique C-terminal lid domain to specifically target host UBC9 and dismantle the SUMOylation machinery, a strategy that is absent in other known bacterial pathogens.

### SseK1 systematically reprograms the host SUMOylome during *Salmonella* **infection**

To gain a comprehensive and unbiased view of the SUMOylation dynamics during *Salmonella* infection, we employed a quantitative proteomic approach utilizing the WaLP-K-ε-GG enrichment strategy^28^, which allows for the site-specific identification of SUMO-modified lysines (Fig. 7a). In STM-infected RAW264.7 macrophages, we identified a total of 303 SUMOylated proteins, with individual targets harboring between one and more than five distinct SUMOylation sites (Fig. 7b and Supplementary Table 2). Functional annotation revealed that these substrates are involved in diverse cellular architectures, such as transcriptional regulation, translational machinery, metabolic pathways, and cytoskeletal organization (Fig. 7c). Strikingly, 95 of these proteins (31%) were not previously documented in the Compendium of Protein Lysine Modifications (CPLM) database, representing novel SUMOylation targets (Fig. 7c). KEGG pathway analysis further enriched 15 of these substrates—including Arf1, Cdc42, Myd88, Ptprc, Rhoa, Tnf, Tuba1b, Tubb2a, Tubb2b, Tubb5, Actb, Actg1, Gapdh, Rps3, and Tubb4b—within the *Salmonella* infection pathway (KEGG mmu05132), suggesting that SUMOylation is a central regulatory node in the host-pathogen interface (Supplementary Fig. 6).

**Fig. 7.**
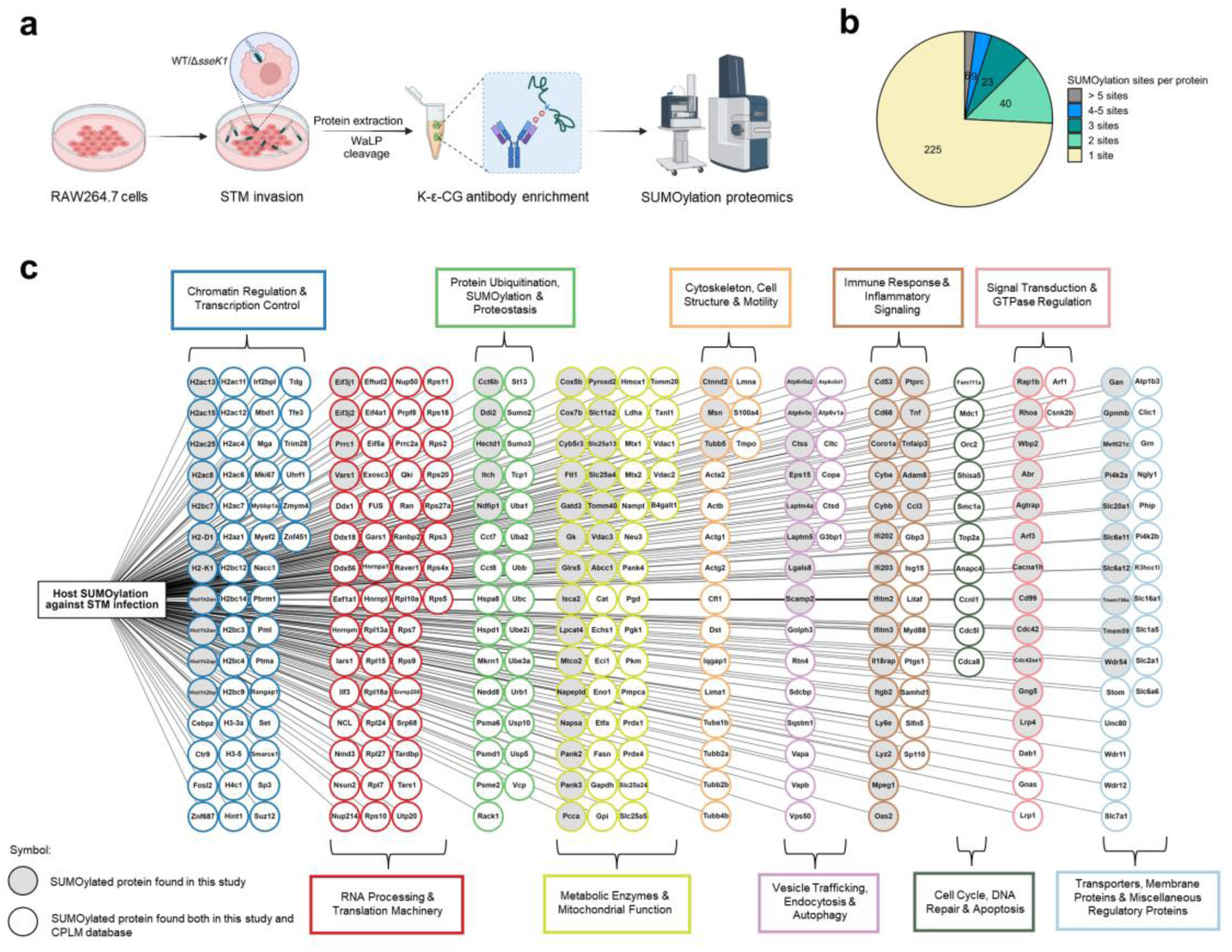
Host SUMOylome during *Salmonella* infection. **a** Schematic of the SUMOylation proteomics workflow. **b** Distribution of SUMOylation site numbers among 303 SUMOylated proteins identified in RAW264.7 cells infected with STM WT and Δ*sseK1* strain (4 hpi). **c** Biological processes associated with the 303 identified SUMOylated proteins.

Given that SseK1 inactivates UBC9, we hypothesized that it would induce a broad reprogramming of the host SUMOylome. By comparing the SUMOylome of cells infected with STM WT versus the Δ*sseK1* mutant, we identified a subset of SUMOylation sites that were specifically suppressed by SseK1 (>3-fold reduction; Fig. 8a). Importantly, the total abundance of these proteins remained stable, indicating that SseK1-mediated effects were specific to the post-translational modification status rather than protein turnover. Gene Ontology (GO) and KEGG enrichment of these SseK1-sensitive targets highlighted key immunological players, such as Myd88, Hspa8, Vcp, Cybb, Isg15, Itch, Pml, Samhd1, Lgals8, and Ptprc (Fig. 8b). We prioritized the core TLR adapter MyD88 and the chaperone Hspa8 for biochemical validation. By incorporating potent deSUMOylation inhibitors during cell lysis, we achieved the first reported detection of SUMO2/3-modified MyD88. IP assays revealed that MyD88 SUMOylation was markedly elevated in Δ*sseK1*-infected cells compared to those infected with STM WT and uninfected (Fig. 8c). A similar trend was observed for Hspa8, where STM WT infection significantly reduced the SUMOylation levels compared to the Δ*sseK1* or uninfected controls (Fig. 8d). These results confirm that SseK1 directly restricts the SUMO2/3 conjugation of critical immune regulators during infection.

**Fig. 8.**
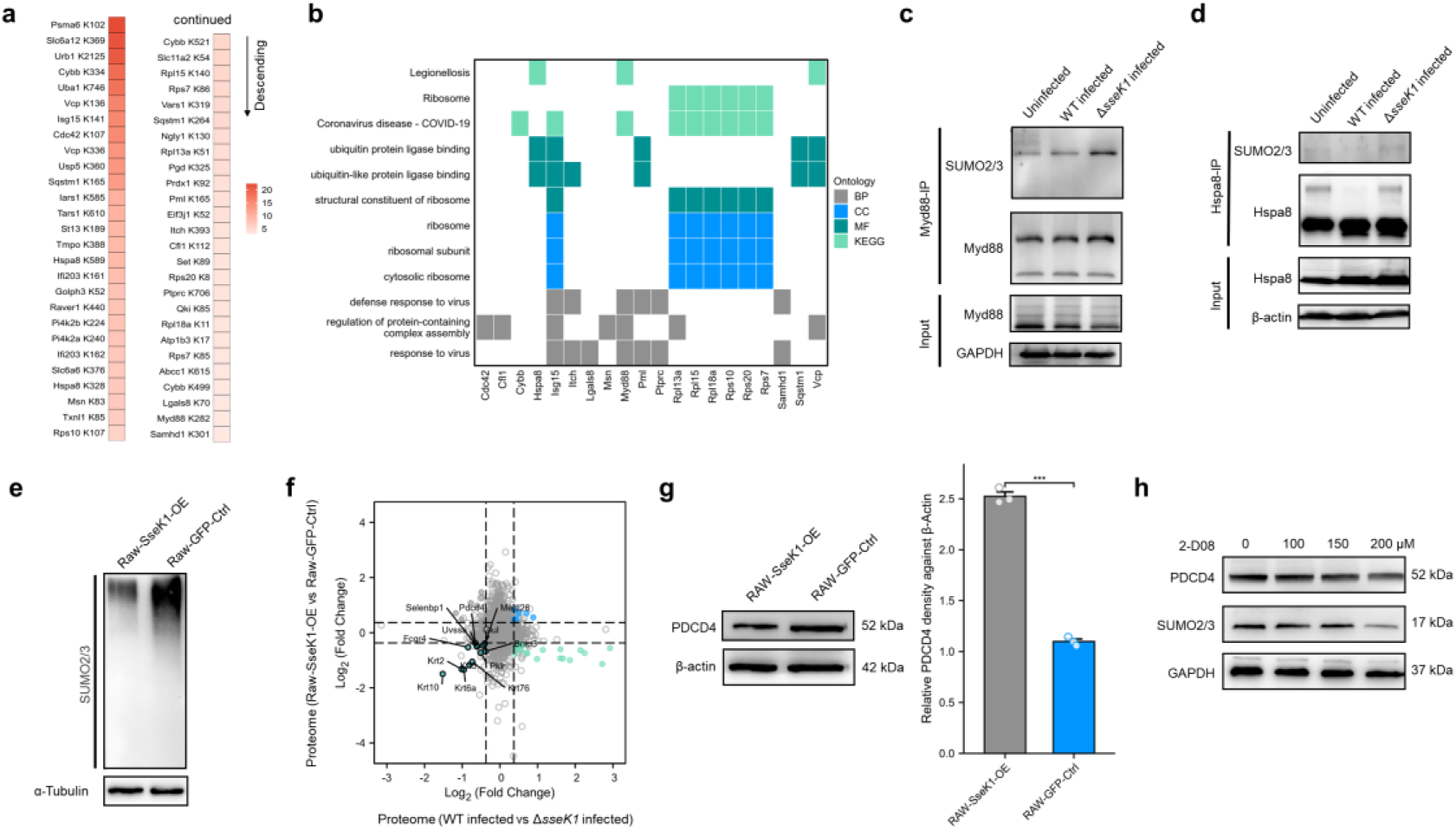
SseK1 reprograms host SUMOylome during *Salmonella* infection. **a** Protein SUMOylation sites showing a >3-fold increase in SUMOylation intensity in STM Δ*sseK1*-infected cells relative to WT. SUMOylation intensity was normalized to the corresponding protein abundance measured in proteomic datasets from STM Δ*sseK1*- or WT-infected RAW264.7 cells. **b** GO and KEGG pathway enrichment analysis of proteins with SUMOylation intensity ratio (Δ*sseK1*/WT) > 3. **c, d** IP analysis of Myd88 and Hspa8 SUMO2/3 modification in RAW264.7 cells infected with STM WT or Δ*sseK1* or left uninfected (4 hpi). **e** Immunoblot analysis of SUMO2/3 modification in RAW-SseK1-OE and RAW-GFP-Ctrl cells. **f** Nine-quadrant plot showing protein abundance changes in two proteomic comparisons (RAW-SseK1-OE vs RAW-GFP-Ctrl and STM WT-infected vs Δ*sseK1*-infected cells). Highlighted proteins are downregulated in both datasets. Fold-change values within dashed lines indicate <|1.3|. **g** Immunoblot analysis of PDCD4 expression in RAW-SseK1-OE and RAW-GFP-Ctrl cells. **h** Immunoblot analysis of PDCD4 and SUMO2/3 levels in RAW264.7 cells treated with increasing concentrations of the SUMOylation inhibitor 2-D08. ***P < 0.001.

Beyond modulating protein activity, SUMOylation is often required for protein conformational stability. To investigate whether SseK1-mediated SUMOylation blockade leads to the degradation of specific host factors, we cross-referenced the proteomes of SseK1-overexpressing macrophages and STM-infected cells. Herein, RAW-SseK1-OE also exhibited reduced SUMO2/3 modification compared with RAW-GFP control cells (Fig. 8e and Supplementary Fig. 7). We identified several proteins whose steady-state levels were significantly depleted in the presence of SseK1, most notably the tumor suppressor and immune regulator PDCD4 (Fig. 8f). Consistent with this, SseK1 overexpression led to a sharp decline in PDCD4 protein levels (Fig. 8g). Furthermore, pharmacological inhibition of the SUMOylation machinery using 2-D08 resulted in a dose-dependent reduction of PDCD4, phenocopying the effect of SseK1 (Fig. 8h). Collectively, these findings demonstrate that SseK1-mediated UBC9 inactivation triggers a global collapse of the host SUMOylome, thereby subverting host defense by altering the activity of adapters like MyD88 and Hspa8 and the stability of effectors like PDCD4.

### SseK1-mediated subversion of host SUMOylation is crucial for *Salmonella* pathogenesis

To determine whether SseK1-mediated subversion of SUMOylation contributes to *Salmonella* pathogenesis, we first evaluated the intracellular fitness of STM in RAW264.7 macrophages. Starting at 2 hpi—a timeframe consistent with the onset of SUMOylation blockade—the Δ*sseK1* mutant exhibited a significant defect in intracellular survival compared to the WT strain (Fig. 9a). To establish a causal link between host SUMOylation and this survival defect, we performed a chemical rescue experiment using the SUMOylation inhibitor 2-D08 (Fig. 9b). While the Δ*sseK1* strain was readily cleared by untreated macrophages, pharmacological inhibition of the host SUMOylation machinery with 2-D08 substantially restored the mutant’s survival to near-WT levels (Fig. 9c). These results indicate that host SUMOylation constitutes a potent antimicrobial defense that *Salmonella* must neutralize via SseK1 to maintain its replicative niche.

**Fig. 9.**
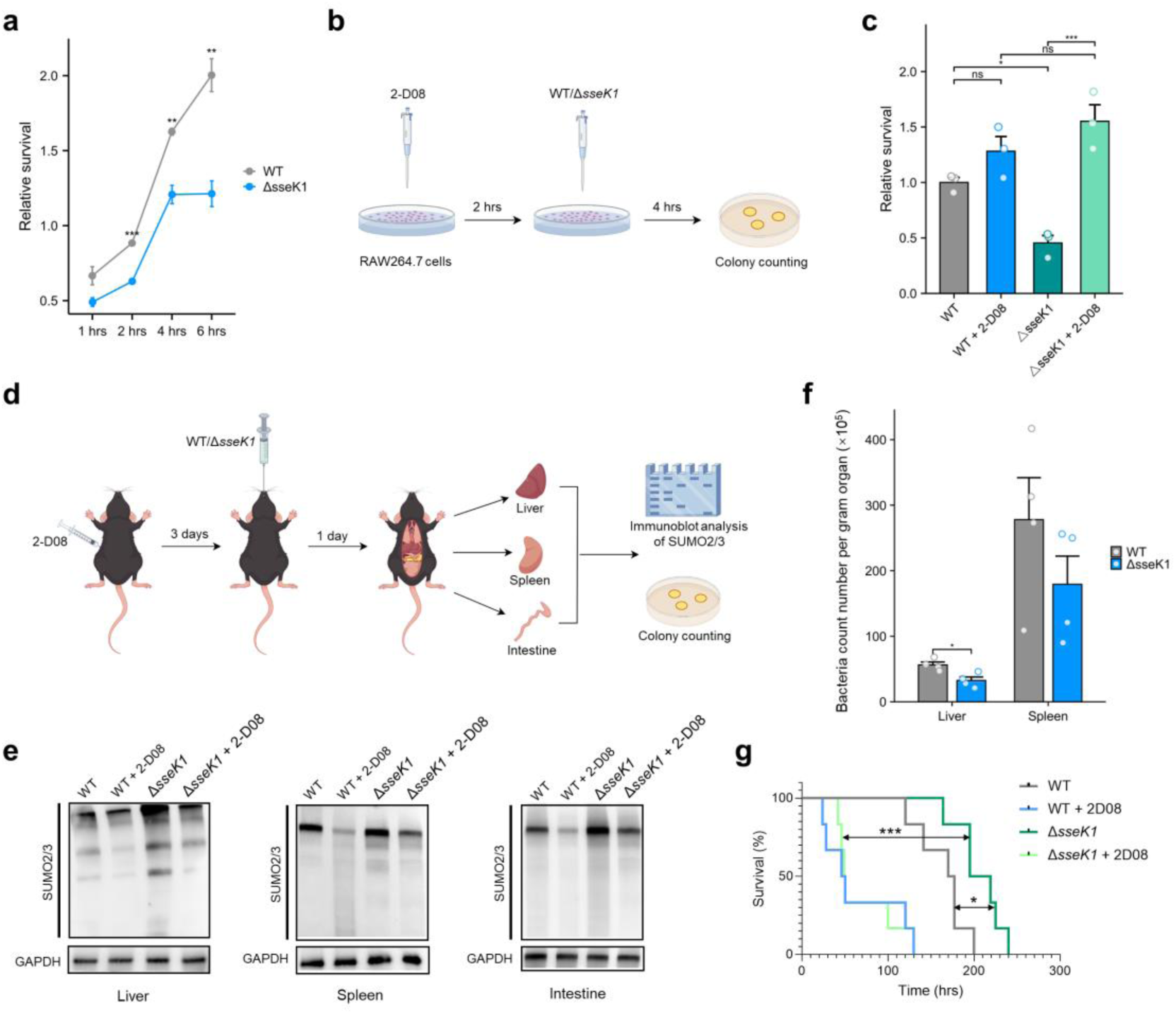
SseK1 enhances *Salmonella* intracellular persistence and virulence by subverting host SUMOylation. **a** Intracellular survival of STM WT and Δ*sseK1* strains in RAW264.7 cells (1-6 hpi). n = 3. **b** Schematic of the experimental workflow for 2-D08 treatment and *Salmonella* infection of RAW264.7 cells. **c** Intracellular survival of STM WT and Δ*sseK1* strains in RAW264.7 cells with or without 2-D08 pretreatment (4 hpi). **d** Schematic of the experimental workflow for 2-D08 administration and STM challenge in C57BL/6 mice. **e** Immunoblot analysis of SUMO2/3 modification in liver, spleen, and intestine tissues from C57BL/6 mice challenged with STM WT or Δ*sseK1*, with or without 2-D08 pretreatment. **f** Bacterial load (CFU per gram of liver or spleen) in C57BL/6 mice infected with STM WT or Δ*sseK1*. n = 4. **g** Survival curves of C57BL/6 mice challenged with STM WT or Δ*sseK1*, with or without 2-D08 pretreatment. n = 6. *P < 0.05; **P < 0.01; ***P < 0.001; ns, not significant.

We further extended these findings to a mouse model of systemic infection (Fig. 9d). Immunoblotting of liver, spleen, and intestinal tissues from infected C57BL/6 mice revealed that Δ*sseK1* infection resulted in markedly higher levels of SUMO2/3 conjugation compared to WT infection. This elevated SUMOylation was successfully suppressed by the administration of 2-D08 (Fig. 9e). Intriguingly, while SseK1 had a negligible effect on SUMO1 modification in macrophage cultures (Supplementary Fig. 5), we observed robust SseK1-dependent suppression of SUMO1 modification in the spleen and intestine *in vivo* (Supplementary Fig. 8). This discrepancy suggests that the substrate specificity or inhibitory breadth of SseK1 may be context-dependent or influenced by tissue-specific host factors. The inability of the Δ*sseK1* mutant to subvert the host SUMOylome translated into significant attenuation *in vivo*. Mice infected with the Δ*sseK1* strain exhibited lower bacterial burdens in both the liver and spleen compared to WT-infected mice (Fig. 9f). Finally, survival analysis demonstrated that the loss of SseK1 significantly attenuated *Salmonella* virulence; however, this attenuation was reversed when host SUMOylation was pharmacologically inhibited with 2-D08 (Fig. 9g). Collectively, these data establish that SseK1-mediated inactivation of the SUMOylation cascade is a pivotal strategy that drives *Salmonella* pathogenesis.

## Discussion

The strategic subversion of host SUMOylation is a hallmark of successful intracellular parasitism^29^. While global suppression of host SUMOylation has been documented across various bacterial taxa, the molecular tactics employed vary significantly. The prevailing paradigm involves the physical depletion of SUMO-cycle enzymes through targeted degradation or proteolytic cleavage. Although *Salmonella* was previously reported to suppress SUMOylation via microRNA-mediated silencing or transcriptional repression, these mechanisms fail to account for the rapid SUMOylation collapse observed within hours of infection. Our study resolves this discrepancy by uncovering a fundamentally different strategy: the enzymatic paralysis of the SUMO E2-conjugating enzyme, UBC9, via SseK1-catalyzed arginine-GlcNAcylation. *Salmonella* utilizes SseK1 as a “surgical strike” tool to rapidly neutralize host SUMOylation-mediated defenses (Fig. 10).

**Fig. 10.**
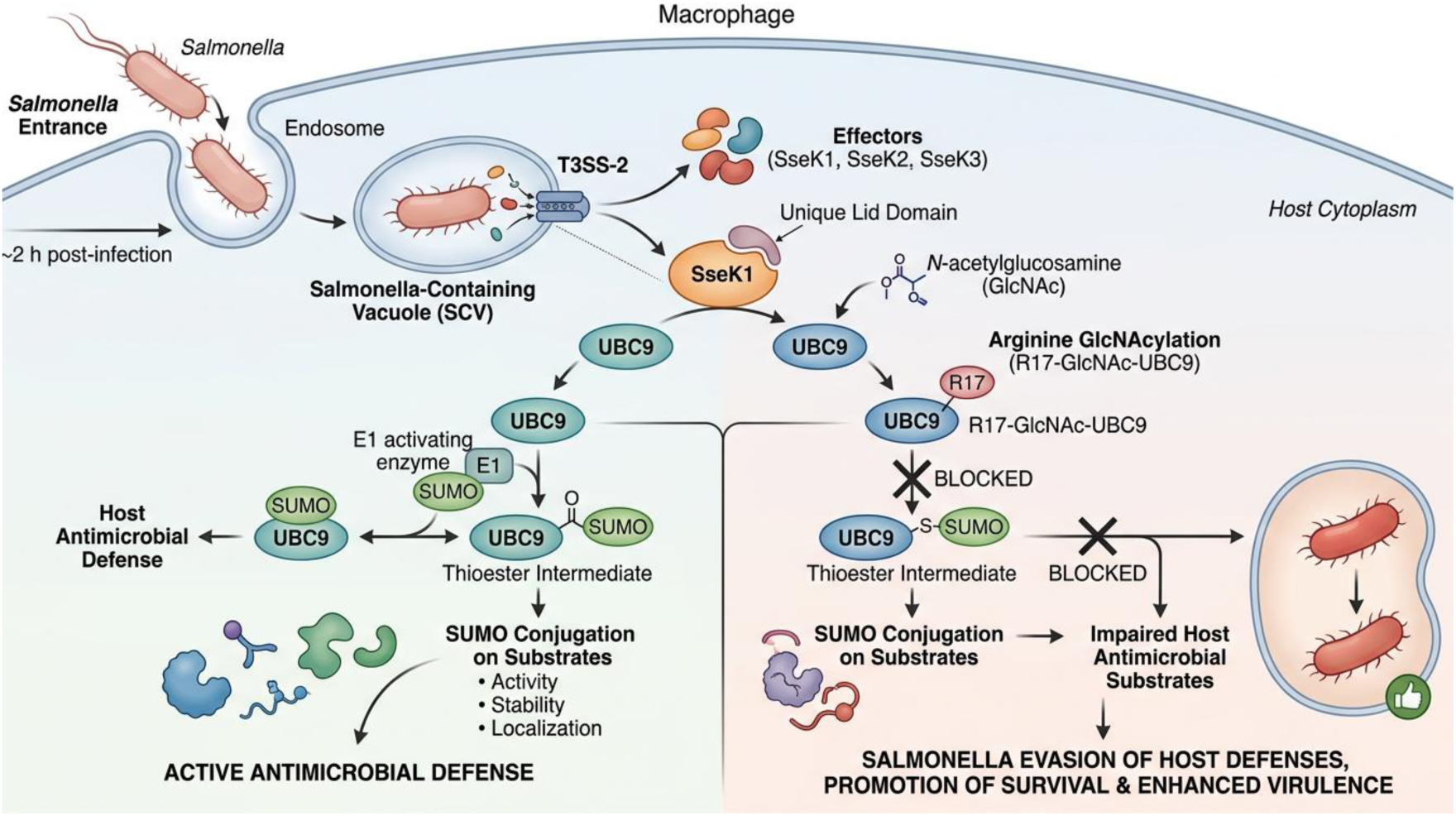
Model of SseK1-mediated subversion of host SUMOylation during *Salmonella* infection. Approximately two hours after entering macrophages, *Salmonella* activates its T3SS-2 to deliver effector proteins, including SseK1, SseK2 and SseK3, into the cytoplasm. Among these effectors, SseK1 uniquely engages UBC9 through its unique lid domain and catalyzes arginine-GlcNAcylation of UBC9 at residue R17. This modification prevents UBC9 from forming the UBC9-SUMO thioester intermediate, thereby blocking SUMO conjugation onto a broad set of antimicrobial substrates and impairing their activity, stability or localization. By blocking this SUMO-dependent arm of the host immune response, *Salmonella* evades host antimicrobial defenses, thereby promoting its survival and enhancing virulence.

The SseK/NleB family utilizes a conserved DXD (Asp-X-Asp) catalytic motif and a lid domain to regulate active-site dynamics, while an HLH (helix-loop-helix) domain facilitates substrate recognition^30, 31^. Notably, some members (SseK1, SseK3, and NleB) undergo auto-GlcNAcylation, a modification that is essential for their ability to inhibit host immune functions^32^. Some members, such as NleB, exhibit broad substrate selectivity toward death domain-containing proteins like TRADD and FADD; this versatility is governed by the second-shell residue Tyr284 adjacent to the catalytic domain, which couples substrate binding with the catalytic process^33^. In contrast, we found that the precision of SseK1 for UBC9 is dictated by its unique C-terminal lid domain, specifically the ARHVQ motif. Structural modeling suggests this domain acts as a molecular clamp, securing UBC9 to facilitate site-specific modification at residue R17. This recruitment mechanism represents a specialized evolutionary innovation exclusive to the *Salmonella* genus, allowing SseK1 to expand its substrate range beyond death domain proteins and effectively exert control over broader host cellular processes.

SUMO-cycle enzymes are governed by a dynamic regulatory network of post-translational modifications (PTMs), including redox states^34^, phosphorylation^35^, ubiquitination^36^, acetylation^37^, and SUMOylation^38^. These modifications function as a sophisticated “switch system” that couples the SUMOylation machinery to the cellular microenvironment^39^. While other pathogens exploit these PTMs to trigger enzyme degradation—such as HPV E6-mediated ubiquitination of UBC9^40^ or Adenovirus Gam1-mediated degradation of the E1 complex^41^—our study provides the first evidence of a bacterial effector utilizing glycosylation to inactivate UBC9. Arginine-GlcNAcylation at R17 represents a highly efficient mode of inactivation. Because R17 is located precisely at the interface where UBC9 non-covalently recruits its SUMO partners, SseK1-mediated modification effectively decouples UBC9 from the SUMO flux without necessitating protein degradation. This “modification-mediated inactivation” may offer distinct advantages over the degradative strategies used by other pathogens. For instance, maintaining stable UBC9 protein levels might prevent host feedback loops that would otherwise sense the loss of the protein and trigger compensatory stress responses^42^. Furthermore, it allows for the potential preservation of non-canonical, SUMOylation-independent functions of UBC9 that might benefit the pathogen’s intracellular lifestyle^43, 44^.

Furthermore, we observed that SseK1 preferentially inhibits SUMO2/3 conjugation in cellular models while exerting a negligible effect on SUMO1. This aligns with previous reports indicating that certain UBC9 inhibitors primarily impair the assembly of poly-SUMO2/3 chains, leaving mono-SUMO1ylation largely unaffected^45^. This preference might reflect an evolutionary adaptation by *Salmonella* to specifically target the stress-inducible SUMO2/3 paralogs, which are more frequently associated with the rapid reprogramming of innate immune signaling than the relatively stable SUMO1 pool^46^. Intriguingly, however, we detected robust SseK1-dependent suppression of SUMO1 modification in the spleen and intestinal tissues of infected mice. This discrepancy suggests that the inhibitory breadth of SseK1 may be context-dependent, potentially modulated by the complex *in vivo* microenvironment—such as localized inflammation or oxidative stress—or by tissue-specific host factors.

SUMOylation serves as a “multipurpose command hub” in antimicrobial defense, coordinating the activity and stability of critical immune regulators. Our quantitative proteomics, providing the first systematic characterization of the host SUMOylome during *Salmonella* infection, identified 303 SUMOylated proteins. The identification of immune adapters like MyD88, Hspa8, and PDCD4 as SseK1-sensitive targets underscores the profound impact of this mechanism. Given that MyD88 SUMOylation may function as a regulatory checkpoint in TLR signaling^47^, its blockade by SseK1 likely dampens inflammatory outputs to favor bacterial survival. Similarly, the reduction of HSPA8 SUMOylation may impair chaperone-mediated autophagy required for bacterial clearance^48^. Furthermore, the SseK1-dependent depletion of PDCD4—which is destabilized upon deSUMOylation^49^—demonstrates how stripping the “SUMO-shield” can indirectly lead to the degradation of essential antimicrobial effectors^50^.

Looking forward, future studies should focus on validating the functional consequences of the novel SUMOylated proteins identified in our SUMOylation proteomics data to further map the host-pathogen interface. Additionally, the discovery of potential “de-arginine-GlcNAcylases” remains a compelling research direction. It is currently unknown if SseK1-mediated modification is reversible, similar to the O-GlcNAcylation system regulated by OGT and OGA^51^. If *Salmonella* possesses such a de-glycosylase, it could provide a “regulatory window” to dynamically tune SUMOylation throughout the infection cycle. Finally, the SseK1-UBC9 axis presents a promising therapeutic target. Given the role of SUMOylation in neurodegenerative diseases^52^, inflammatory disorders^53^, and cancers^54^, the precision with which SseK1 regulates UBC9 activity could be exploited for the development of innovative mRNA-based or enzymatic modulation strategies.

In conclusion, our study defines the SseK1-UBC9 axis as a critical battleground in host-pathogen interactions. By targeting the R17 residue—the central hub for SUMO recruitment—*Salmonella* achieves a systemic collapse of the host SUMOylome. This modification-mediated inactivation strategy not only provides a highly efficient means of subverting immunity but also highlights the remarkable functional complexity evolved by a single bacterial effector.

## Methods

### Bacterial strains and plasmids

*Salmonella enterica* serovar Typhimurium ATCC 14028S and its derivatives were cultured in Luria-Bertani (LB) medium at 37°C. *E. coli* BL21(DE3) was used for recombinant protein expression and purification. Antibiotics were added as required for plasmid maintenance. All bacterial strains and plasmids used in this study are listed in Supplementary Table 3.

### Cell lines and culture conditions

RAW264.7 macrophages, RAW264.7 cells stably expressing SseK1 (RAW-SseK1-OE), RAW264.7 cells expressing GFP (RAW-GFP-Ctrl), and HEK293T cells were cultured in Dulbecco’s modified Eagle medium (DMEM, Gibco) supplemented with 10% fetal bovine serum (FBS), 100 U/mL penicillin, and 100 μg/mL streptomycin at 37°C in 5% CO₂. RAW-SseK1-OE and RAW-GFP-Ctrl cell lines were generated by Ubigene Biosciences (Guangzhou, China).

### Animals

Female C57BL/6J mice (6-8 weeks old) were purchased from Pengyue Laboratory Animal Technology Co., Ltd (Jinan, China), randomly assigned to experimental groups, and housed under standardized conditions with food and water ad libitum. All animal procedures complied with institutional and national guidelines for the Care and Use of Laboratory Animals and were approved by the Animal Ethics Committee of Shandong First Medical University (Permit No. W202103170256).

### Cell infection models

STM 14028S strains were grown overnight in LB at 37°C with shaking, diluted 1:100 into fresh LB, and cultured to an OD600 of 0.5. For SseK1-complemented strains (harboring pBAD24 vectors), 0.2% L-arabinose was added at OD600 = 0.5 and incubation continued for an additional 2 hrs. RAW264.7 or HEK293T cells were seeded into 24- (5 × 10⁵ cells per well) or 96- (5 × 10⁵ cells per well) well plates and infected at an MOI of 20. At 1 hpi, cells were washed twice with PBS and incubated with DMEM containing 100 μg/mL gentamicin to eliminate extracellular bacteria. Cells before infection served as the 0 hr control. For 2-D08 pretreatment, 50 μM 2-D08 was added 2 hrs prior to infection.

### Mouse infection models

After 1 week of acclimatization, C57BL/6J mice were randomly assigned to experimental groups (n = 4 or 6). Mice were orally inoculated with 2 × 10⁸ CFU of STM WT or Δ*sseK1*. For 2-D08 treatment, 5 mg/kg 2-D08 was administered intraperitoneally 3 days before infection. For immunoblot or CFU assays, mice were euthanized at 1 dpi, and liver, spleen, and intestine tissues were collected. For survival assays, mouse survival was monitored every 1 hr and used to generate survival curves.

### Real-time quantitative PCR

At designated time points following infection, cells were lysed in 1% Triton X-100, and total RNA was extracted using the RNAprep Pure Cell/Bacteria Kit (TIANGEN). The cDNA synthesis was performed using the HiScript III 1st Strand cDNA Synthesis Kit (Vazyme). qPCR reactions were prepared with TB Green Premix Ex Taq II (TaKaRa) and analyzed using an Applied Biosystems 7500 system. UBC9 expression was quantified using the 2-ΔΔCT method with 16S rRNA as the internal reference.

### Immunofluorescence assay

At designated time points, infected cells were fixed with 4% paraformaldehyde (PFA) for 30 min and permeabilized with 0.25% Triton X-100 for 10 min. After blocking with 5% BSA for 1 hr, cells were incubated with primary antibodies (SUMO1 mouse mAb, 67559-1-Ig; SUMO2/3 rabbit pAb, 11251-1-AP; Proteintech) for 2 hrs at room temperature, followed by Alexa Fluor 488- or 594-conjugated secondary antibodies. Images were acquired with a confocal laser scanning microscope (LSM900, Carl Zeiss), and fluorescence intensity was quantified using ImageJ. For high-content screening, images were obtained using the ImageXpress Micro system (Molecular Devices).

### Colony-forming unit (CFU) assay

Infected cells or mouse tissues were collected at designated time points and lysed in PBS containing 1% Triton X-100. Lysates were serially diluted and plated on LB agar, incubated overnight at 37°C, and CFUs counted. Relative survival was normalized to STM WT infection or the 0 hr time point.

### Transcriptomic, proteomic, and SUMOylation proteomic assays

Cells from infection models or from RAW-SseK1-OE and RAW-GFP-Ctrl cultures were harvested and flash-frozen in liquid nitrogen. Transcriptomic sequencing was performed using the Illumina NovaSeq 6000 platform (Novogene, Beijing). Proteomic analysis was performed using Q Exactive HF-X (Thermo Fisher) or timsTOF Pro 2 (Bruker) mass spectrometers (PTM BIO, Hangzhou). For SUMOylation proteomics, cell pellets were lysed in RIPA buffer containing 1% PMSF and 20 mM N-ethylmaleimide (to stabilize SUMO conjugates), digested with trypsin, and centrifuged. Supernatants were incubated with anti-diglycine lysine antibody-conjugated agarose beads (PTM BIO) at 4°C overnight. Beads were sequentially washed with IP buffer and deionized water, and bound peptides were eluted with 0.1% trifluoroacetic acid (three times), dried, desalted with C18 ZipTips, dried again, separated by Vanquish Neo UHPLC, and analyzed using an Orbitrap Astral mass spectrometer (Thermo Fisher). SUMOylation proteomics were performed by PTM BIO Co., Ltd. (Hangzhou, China).

### Y2H screening and validation

SseK1 was cloned into pGBKT7 and co-transformed with a pGADT7-RAW264.7 cDNA library into Y2H Gold yeast cells. Positive clones on SD/-Leu/-Trp/-His/X-α-gal plates (DDO/X/A) were transferred to SD/-Leu/-Trp/-His/-Ade/X-α-gal/AbA plates (QDO/X/A) for confirmation and subsequently sequenced. The SseK1-UBC9 interaction was validated by co-transforming pGBKT7-SseK1 and pGADT7-UBC9. pGBKT7-53/pGADT7-T and pGBKT7-Lam/pGADT7-T served as positive and negative controls, respectively. All Y2H assays were performed by OE Biotech Co., Ltd (Shanghai, China).

### Recombinant protein expression and purification

Recombinant plasmids were transformed into *E. coli* BL21(DE3) for expression of SseK1, SseK2, SseK3 and their mutants, UBC9, Arg-GlcNAcylated UBC9, SUMO1, and SUMO2. Protein expression was induced with 0.1 mM IPTG at OD600 = 0.4-0.6, followed by overnight incubation at 16°C. Cells were collected, lysed, and proteins purified using Ni²⁺-NTA affinity chromatography. When required, the N-terminal 6×His tag was removed using PPase (pGL01 vector system), and proteins were further purified by size-exclusion chromatography (Superdex 200 Increase Columns)

### Pull-down assay

His-tagged SseK1 was used as bait and tag-free UBC9 (after PPase treatment) as prey. Proteins were incubated with Ni²⁺-NTA resin, washed, and analyzed by SDS-PAGE. Controls consisted of bait or prey alone.

### BLI assay

BLI was performed using an Octet RED96 system (Sartorius) at 25°C. UBC9 was desalted using a Mini Trap G-25 column (Cytiva), biotinylated with EZ-Link NHS-Biotin (Thermo Fisher) for 30 min at room temperature, and desalted again. SseK1 was desalted and adjusted to 1 μM. Streptavidin (SA) biosensors were equilibrated in PBS, loaded with biotinylated UBC9 for 240 s, equilibrated again, and then dipped into SseK1 solution for 420 s. Dissociation was measured in PBS. Binding curves and Kd values were obtained using RED96 software.

### MST assay

SUMO1 or SUMO2 was desalted and labeled with the RED-NHS Protein Labeling Kit (NanoTemper). UBC9 or Arg-GlcNAcylated UBC9 was desalted and serially diluted (5 μM to 0.15 nM), mixed with 200 nM labeled SUMO proteins, loaded into Monolith NT.115 capillaries, and measured at 25°C (40% LED, medium MST power). Data was analyzed using MO.Affinity Analysis software to determine Kd values.

### *In vitro* arginine-GlcNAcylation reaction

The 50 µL reactions containing 1 μg SseK proteins or mutants, 5 μg UBC9, 1 mM UDP-GlcNAc, 2 mM MnCl₂, 20 mM Tris-HCl (pH 8.5), and 150 mM NaCl were incubated at 37°C for 2 hrs. Reactions were terminated by boiling at 95°C for 5 min and analyzed by SDS-PAGE and immunoblotting.

### IP and Co-IP

For Flag-IP, HEK293T cells were transfected with pCDNA3.1-3×Flag-UBC9, pEGFP-N1-SseK1, and/or control plasmids using Lipofectamine 2000. At 20 hpi, cells were infected with STM WT or Δ*sseK1* as described, lysed in RIPA buffer with 1% PMSF, and incubated with anti-Flag magnetic beads (Selleck) overnight at 4°C. For Myd88-IP and HSPA8-IP, RAW264.7 cells were infected for 4 hrs, lysed in RIPA buffer containing 1% PMSF and 20 mM N-ethylmaleimide, and incubated with Myd88 or HSPA8 antibodies overnight at 4°C, followed by protein A/G bead incubation for 3 hrs. Beads were washed 4-6 times with PBST, boiled in SDS loading buffer, and analyzed by immunoblotting.

### Immunoblot assay

Standard immunoblotting procedures were used. Extracts were prepared in RIPA buffer (with PMSF and N-ethylmaleimide when required), separated using 12% SDS-PAGE, and transferred to nitrocellulose membranes (300 mA, 1 hr). Membranes were blocked in 5% milk, incubated with primary antibodies overnight at 4°C and with secondary antibodies for 1 hr at room temperature. Bands were visualized by ECL and quantified using ImageJ. Unique primary antibodies used in this study included UBC9 (CST, #4786), SUMO1 (Proteintech, 67557-1-lg), SUMO2/3 (Proteintech, 67154-1-lg), arginine-GlcNAcylation antibody (Abcam, EPR18251), Myd88 (Proteintech, 67969-1-lg), HSPA8 (Proteintech, 10654-1-AP), PDCD4 (Proteintech, 84162-3-RR).

### Identification of arginine-GlcNAcylation sites

Putative Arg-GlcNAcylated UBC9 bands were subjected to in-gel tryptic digestion. Peptides were separated using an EASY-nLC 1200 system, ionized by NSI, and analyzed on an Orbitrap Exploris 480 mass spectrometer. Data were searched using Proteome Discoverer 2.4 with arginine GlcNAcylation specified as a variable modification. Peptides required scores >20 and high confidence. Analyses were performed by PTM BIO.

### Protein modeling and visualization

Amino acid sequences of SseK1, SseK2, SseK3, UBC9, and SUMO2 were retrieved from UniProt. Protein or protein-complex structures (SseK1, SseK2, SseK3, SseK1-UBC9, SUMO2-UBC9) were predicted using AlphaFold3 (https://alphafold3.org/) with Model 0 selected for analysis. Sequence alignment and structural visualization were performed using PyMOL.

### Statistical analysis

Data are presented as mean ± SEM. For two-group comparisons, Student’s t-test, Welch’s t-test, or the Wilcoxon rank-sum test was used depending on normality and variance. For multiple-group comparisons, one-way ANOVA, Welch’s ANOVA, or the Kruskal-Wallis test was applied as appropriate. For survival curves, Mantal-Cox test was used for two-group comparison. Statistical methods were selected according to data distribution characteristics and performed using R packages (stats 4.2.1, car 3.1-0). Significance thresholds were set at P < 0.05, **P < 0.01, ***P < 0.001, and ns (not significant). Exact n values (representing independent experiments or animals) are indicated in the figure legends.

## Author contributions

B.Q.L. conceived and directed the project. B.Q.L., Z.R.M., and H.J.Z. designed the experiments and interpreted the data. Z.R.M., Y.Y.Z., H.J.Z. performed most of the experiments. Y.H.Z. and X.Y.W. conducted additional experiments and contributed to data interpretation. B.Q.L. supervised experiments and assisted with data interpretation. B.Q.L., Z.R.M., and H.J.Z. wrote the manuscript with input from all authors.

## Acknowledgements

This work was supported by National Natural Science Foundation of China [3257206, 32170034, 32000647], the Taishan Scholar Project of Shandong Province [tsqn202211216], the Natural Science Foundation of Shandong Province [ZR2023YQ060, ZR2024QC359], the Joint Innovation Team for Clinical & Basic Research of Shandong First Medical University [202410], Key Project in Medical and Health Technology of Shandong Province [202401060550].

## References

1. Liu J, Qian C, Cao X. Post-Translational Modification Control of Innate Immunity. Immunity 45, 15–30 (2016).

2. Vertegaal ACO. Signalling mechanisms and cellular functions of SUMO. Nature reviews Molecular cell biology 23, 715–731 (2022).

3. Tharuka MDN, Courelli AS, Chen Y. Immune regulation by the SUMO family. Nature reviews Immunology 25, 608–620 (2025).

4. Goffeney A, et al. SUMO operates from a unique long tandem repeat to keep innate immunity in check. Nucleic acids research 53, (2025).

5. Cai J, et al. SUMOylation protects against sepsis-associated acute kidney injury by stabilizing IkappaBalpha. Molecular therapy : the journal of the American Society of Gene Therapy, (2025).

6. Gu A, et al. SENP6 Restrains NLRP3 Inflammasome Activation via DeSUMOylation-Driven K48-Linked Ubiquitination of NLRP3 in Acute Lung Injury. Research (Washington, DC) 9, 1069 (2026).

7. Garvin AJ, et al. SUMO4 promotes SUMO deconjugation required for DNA double-strand-break repair. Molecular cell 85, 877–893.e879 (2025).

8. He Q, Chen J, Chen S, Gao X. SUMOylation modulates the dual functions of Kruppel homolog 1 in transcriptional regulation of Broad-Complex expression. Journal of advanced research 75, 123–135 (2025).

9. Domingues P, et al. Global Reprogramming of Host SUMOylation during Influenza Virus Infection. Cell reports 13, 1467–1480 (2015).

10. Fritah S, et al. Sumoylation controls host anti-bacterial response to the gut invasive pathogen Shigella flexneri. EMBO reports 15, 965–972 (2014).

11. Zhu G, Tong N, Zhu Y, Wang L, Wang Q. The crosstalk between SUMOylation and immune system in host-pathogen interactions. Critical reviews in microbiology 51, 164–186 (2025).

12. Ma X, Zhao C, Xu Y, Zhang H. Roles of host SUMOylation in bacterial pathogenesis. Infection and immunity 91, e0028323 (2023).

13. Xu H, et al. Multiple Enzymes Expressed by the Gut Microbiota Can Transform Typhaneoside and Are Associated with Improving Hyperlipidemia. Advanced science (Weinheim, Baden-Wurttemberg, Germany) 12, e2411770 (2025).

14. Ribet D, et al. Listeria monocytogenes impairs SUMOylation for efficient infection. Nature 464, 1192–1195 (2010).

15. Loison L, et al. Staphylococcus warneri dampens SUMOylation and promotes intestinal inflammation. Gut microbes 17, 2446392 (2025).

16. Lapaquette P, et al. Shigella entry unveils a calcium/calpain-dependent mechanism for inhibiting sumoylation. eLife 6, (2017).

17. Li Y, Perez-Gil J, Lois LM, Varejao N, Reverter D. Broad-spectrum ubiquitin/ubiquitin-like deconjugation activity of the rhizobial effector NopD from Bradyrhizobium (sp. XS1150). Communications biology 7, 644 (2024).

18. Sa-Pessoa J, et al. Klebsiella pneumoniae Reduces SUMOylation To Limit Host Defense Responses. mBio 11, (2020).

19. Verma S, et al. Salmonella Engages Host MicroRNAs To Modulate SUMOylation: a New Arsenal for Intracellular Survival. Molecular and cellular biology 35, 2932–2946 (2015).

20. Kumar P, et al. Bidirectional regulation between AP-1 and SUMOylation pathway genes modulates inflammatory signaling during Salmonella infection. Journal of cell science 135, (2022).

21. Newson JPM, et al. Salmonella Effectors SseK1 and SseK3 Target Death Domain Proteins in the TNF and TRAIL Signaling Pathways. Molecular & cellular proteomics : MCP 18, 1138–1156 (2019).

22. Lu X, et al. Effects of Salmonella enterica serovar typhimurium sseK1 on macrophage inflammation-related cytokines and glycolysis. Cytokine 140, 155424 (2021).

23. Gunster RA, Matthews SA, Holden DW, Thurston TLM. SseK1 and SseK3 Type III Secretion System Effectors Inhibit NF-kappaB Signaling and Necroptotic Cell Death in Salmonella-Infected Macrophages. Infection and immunity 85, (2017).

24. Wang KC, Huang CH, Huang CJ, Fang SB. Impacts of Salmonella enterica Serovar Typhimurium and Its speG Gene on the Transcriptomes of In Vitro M Cells and Caco-2 Cells. PloS one 11, e0153444 (2016).

25. Zhang K, et al. Age-dependent enterocyte invasion and microcolony formation by Salmonella. PLoS pathogens 10, e1004385 (2014).

26. Rodenburg W, et al. Salmonella induces prominent gene expression in the rat colon. BMC microbiology 7, 84 (2007).

27. Knipscheer P, van Dijk WJ, Olsen JV, Mann M, Sixma TK. Noncovalent interaction between Ubc9 and SUMO promotes SUMO chain formation. The EMBO journal 26, 2797–2807 (2007).

28. Lumpkin RJ, et al. Site-specific identification and quantitation of endogenous SUMO modifications under native conditions. Nature communications 8, 1171 (2017).

29. Xu Y, et al. Host SUMOylation in bacterial infections and immune defense mechanisms. Front Microbiol 16, 1621137 (2025).

30. Park JB, et al. Structural basis for arginine glycosylation of host substrates by bacterial effector proteins. Nature communications 9, 4283 (2018).

31. Esposito D, et al. Structural basis for the glycosyltransferase activity of the Salmonella effector SseK3. J Biol Chem 293, 5064–5078 (2018).

32. Xue J, et al. Auto Arginine-GlcNAcylation Is Crucial for Bacterial Pathogens in Regulating Host Cell Death. Front Cell Infect Microbiol 10, 197 (2020).

33. Garcia-Garcia A, et al. NleB/SseK-catalyzed arginine-glycosylation and enteropathogen virulence are finely tuned by a single variable position contiguous to the catalytic machinery. Chem Sci 12, 12181–12191 (2021).

34. Bossis G, Melchior F. Regulation of SUMOylation by reversible oxidation of SUMO conjugating enzymes. Mol Cell 21, 349–357 (2006).

35. Su YF, Yang T, Huang H, Liu LF, Hwang J. Phosphorylation of Ubc9 by Cdk1 enhances SUMOylation activity. PloS one 7, e34250 (2012).

36. Itahana Y, Yeh ET, Zhang Y. Nucleocytoplasmic shuttling modulates activity and ubiquitination-dependent turnover of SUMO-specific protease 2. Mol Cell Biol 26, 4675–4689 (2006).

37. Hsieh YL, et al. Ubc9 acetylation modulates distinct SUMO target modification and hypoxia response. Embo j 32, 791–804 (2013).

38. Knipscheer P, et al. Ubc9 sumoylation regulates SUMO target discrimination. Mol Cell 31, 371–382 (2008).

39. Chang HM, Yeh ETH. SUMO: From Bench to Bedside. Physiol Rev 100, 1599–1619 (2020).

40. Heaton PR, Deyrieux AF, Bian XL, Wilson VG. HPV E6 proteins target Ubc9, the SUMO conjugating enzyme. Virus Res 158, 199–208 (2011).

41. Boggio R, Colombo R, Hay RT, Draetta GF, Chiocca S. A mechanism for inhibiting the SUMO pathway. Mol Cell 16, 549–561 (2004).

42. Costa-Mattioli M, Walter P. The integrated stress response: From mechanism to disease. Science 368, (2020).

43. Wu H, et al. UBC9 ameliorates diabetic cardiomyopathy by modulating cardiomyocyte mitophagy through NEDD4/RUNX2/PSEN2 axis. Metabolism 168, 156264 (2025).

44. Tang B, et al. UBC9 regulates cardiac sodium channel Na(v)1.5 ubiquitination, degradation and sodium current density. J Mol Cell Cardiol 129, 79–91 (2019).

45. Wiechmann S, et al. Site-specific inhibition of the small ubiquitin-like modifier (SUMO)-conjugating enzyme Ubc9 selectively impairs SUMO chain formation. J Biol Chem 292, 15340–15351 (2017).

46. Acuna ML, et al. Alternative splicing of the SUMO1/2/3 transcripts affects cellular SUMOylation and produces functionally distinct SUMO protein isoforms. Scientific reports 13, 2309 (2023).

47. Li Q, et al. SPOP promotes ubiquitination and degradation of MyD88 to suppress the innate immune response. PLoS Pathog 16, e1008188 (2020).

48. Miao C, et al. HSPA8 regulates anti-bacterial autophagy through liquid-liquid phase separation. Autophagy 19, 2702–2718 (2023).

49. Wang YX, et al. Downregulation of PDCD4 by deSUMOylation associates with the progression of gestational trophoblastic disease. Placenta 130, 17–24 (2022).

50. Zhang X, Zhang J, Li F, Luo Y, Jiang S. PDCD4-mediated downregulation of Listeria monocytogenes burden in macrophages. Central-European journal of immunology 46, 38–46 (2021).

51. Lu P, et al. Cryo-EM structure of human O-GlcNAcylation enzyme pair OGT-OGA complex. Nat Commun 14, 6952 (2023).

52. Wan L, et al. Age-related p53 SUMOylation accelerates senescence and tau pathology in Alzheimer’s disease. Cell Death Differ 32, 837–854 (2025).

53. Youssef A, et al. Vagal stimulation ameliorates murine colitis by regulating SUMOylation. Sci Transl Med 16, eadl2184 (2024).

54. Seeler JS, Dejean A. SUMO and the robustness of cancer. Nat Rev Cancer 17, 184–197 (2017).

